# Impact of human airway epithelial cellular composition on SARS-CoV-2 infection biology

**DOI:** 10.1101/2021.07.21.453304

**Authors:** Ying Wang, Melissa Thaler, Dennis K. Ninaber, Anne M. van der Does, Natacha S. Ogando, Hendrik Beckert, Christian Taube, Clarisse Salgado-Benvindo, Eric J. Snijder, Peter J. Bredenbeek, Pieter S. Hiemstra, Martijn J. van Hemert

**Affiliations:** Department of Pulmonology, Leiden University Medical Center, Leiden, The Netherlands; Department of Medical Microbiology, Leiden University Medical Center, Leiden, The Netherlands; Department of Pulmonary Medicine, University Medical Center Essen – Ruhrlandklinik, Essen, Germany

**Keywords:** SARS-CoV-2, human airway epithelial cells, infection biology, differentiation, cellular composition

## Abstract

Infection biology and pathogenesis of severe acute respiratory syndrome coronavirus 2 (SARS-CoV-2), the causative agent of the coronavirus disease 2019 (COVID-19), are incompletely understood. Here, we assessed the impact of airway epithelial cellular composition on infection in air-liquid interface (ALI) cultures of differentiated primary human tracheal (PTEC) and bronchial epithelial cells (PBEC). We first compared SARS-CoV-2 infection kinetics, related antiviral and inflammatory responses, and viral entry factors in PTEC and PBEC. Next, the contribution of differentiation time was investigated by differentiating ALI-PTEC/PBEC for 3-5 weeks and comparing dynamics of viral replication/spread, cellular composition and epithelial responses. We observed a gradual increase in viral load with prolonged culture duration. Ciliated and goblet cells were predominantly infected in both PTEC and PBEC. Immunofluorescence analysis and RT-qPCR showed that compared to other cell types mainly ciliated and goblet cell numbers were affected by increased culture duration. An increased proportion of these two target cell types was associated with increased viral load. Furthermore, modulation of cellular composition using IL-13 and the Notch signaling inhibitor DAPT, underlined the importance of both ciliated and goblet cells for infection. DAPT treatment resulted in a lower viral load and a relative increase in ciliated cells at the expense of goblet cells, compared to IL-13 treated cultures in which both cell types were present and viral load was higher.

In conclusion, our results identify cellular composition as a contributing factor to airway epithelial susceptibility to SARS-CoV-2.

**IMPORTANCE:** In this study, we determined an effect of culture duration and airway cellular composition of ALI-PBEC and ALI-PTEC cultures on SARS-CoV-2 infection. We found that SARS-CoV-2 infection was increased with prolonged cell culture time and the total percentage and proportion of ciliated and goblet cells played an important role in infection level, suggesting that airway epithelial differentiation/maturation levels may in part determine susceptibility of SARS-CoV-2 infection.

The development of effective therapies either targeting virus replication or pathogenesis against SARS-CoV-2 requires robust cell culture-based infection models to test small molecules and biologicals. Therefore, it is important to identify factors that are essential for reliably modeling SARS-CoV-2-airway epithelial cell interactions. This study sheds light on virus-airway epithelial cell interactions and adds to the complexity of SARS-CoV-2 cell tropism in the airways. In addition, the effect of IL-13 on viral infection hints at a causal connection between SARS-CoV-2 infection and (allergic) asthma.

## INTRODUCTION

Since December 2019, severe acute respiratory syndrome coronavirus 2 (SARS-CoV-2) has rapidly spread worldwide. The burden of the associated coronavirus disease 2019 (COVID-19) on human health has an enormous social and economic impact (1). In an immense effort to combat COVID-19, several successful vaccines have been developed and approved for large scale vaccination campaigns across the globe. However, there is not yet a specific antiviral drug that was proven to be beneficial for relevant patient outcomes. Furthermore, the emergence of virus variants, such as those first observed in the United Kingdom, South Africa, Brazil and India (2–5), which are associated with a rapid increase in COVID-19 cases (6, 7), illustrates the threat of these viruses to prolong the current pandemic or lead to new large outbreaks in the future. These recent developments also underscore the need to develop therapeutic options, which requires a better understanding of SARS-CoV-2 infection biology and pathogenesis.

SARS-CoV-2 was first isolated from the lower respiratory tract of patients presenting with COVID-19 (8, 9). The respiratory tract spans from the nasal cavity to the terminal bronchioles. It is lined by the airway epithelium, which includes many cell types: mainly ciliated, goblet, club and basal cells (10, 11). The epithelium serves as first barrier to SARS-CoV-2 infection and therefore epithelial immune responses (antiviral responses, inflammatory responses) play an important role in the outcome of infection. *In vitro* cultures of Vero E6 cells, lung epithelial cell lines like Calu-3 cells, and differentiated primary human bronchial epithelial cells have enabled studies on the kinetics of SARS-CoV-2 infection in more detail (12, 13). Recent reports showed that SARS-CoV-2 targets ciliated and secretory cells (14, 15), or ciliated cells with strands of mucus attached to the cilia tips (16). SARS-CoV-2 cellular tropism depends on host proteins that are involved in virus entry, including angiotensin converting enzyme 2 (ACE2) and proteases like transmembrane protease serine 2 (TMPRSS2), as well as alternative receptors (cluster of differentiation 147 [CD147], 78-kDa glucose-regulated protein [GRP78], tyrosine-protein kinase receptor UFO [AXL]), which all have been demonstrated to be present in human airway epithelial cells (17–20). The involvement of these and other factors in SARS-CoV-2 infection biology and pathogenesis can be therefore studied in an *in vitro* model of human primary airway epithelial cells. However, the airway epithelium is a dynamic system with cellular compositions that vary throughout the respiratory tract (10) and that is in constant turnover (21). In addition, in many patients with lung diseases such as asthma or chronic obstructive pulmonary disease (COPD), epithelial cellular composition is altered (11). Cellular composition also can be modulated by other factors (e.g. the Notch signaling inhibitor DAPT and the Th2 cytokine IL-13) *in vitro* (22–24). Since SARS-CoV-2 was shown to target mainly ciliated and goblet cells, we hypothesized that cellular composition of the airway epithelium affects SARS-CoV-2 infection and this could contribute to regional and airway disease-associated differences in infection biology as observed in human lungs.

To this end, we used primary human bronchial (PBEC) or tracheal epithelial cells (PTEC) and differentiated them by culture at the air-liquid interface (ALI) to generate the cell types that constitute the airway epithelium. We characterized virus replication, spread, localization, immune responses, and expression of SARS-CoV-2-entry factors, as well as compared cellular composition between both types of cultures. Furthermore, we investigated how differences in culture and infection characteristics were influenced by the duration of culturing and by modulation of the cellular composition using DAPT and IL-13.

## RESULTS

### SARS-CoV-2 efficiently infects differentiated human primary bronchial epithelial cells

To investigate the effect of differences in cellular composition on SARS-CoV-2 infection of airway epithelial cells, we first aimed to establish a reliable infection model. Therefore, mixes of ALI-PBEC from 4 or 5 donors were differentiated for 5 weeks as previously reported (14). By using mixes of primary cells from several donors (donor mix) for optimization of our model, we aimed to limit the impact of possible inter-donor variation, while maintaining the characteristics of using primary cells. We decided to perform infections at a relatively low multiplicity of infection (MOI) to initially only infect a fraction of susceptible cells and observe the virus spread across the epithelium through time. Four independent experiments were performed using ALI-PBEC derived from the same donor mix. After infection with SARS-CoV-2 (30,000 PFU per insert), we observed an increase in viral load (extracellular copies of viral RNA) over time, resulting in about 10^11^ copies/ml at 72 hours post infection (hpi) (Fig. 1A). Immunofluorescence staining of the epithelial cultures for viral nucleocapsid protein also showed an increase in the number of infected cells over time (Fig. 1B). We next compared infection kinetics in the donor mix and in cultures of the 4 individual donors that were used to establish the mix. Single donor and mixed donor cultures showed comparable infection kinetics (Fig. 1C), and immunofluorescence staining also showed similar numbers of infected cells at 72 hpi (Fig. 1D) thereby supporting the use of these donor mixes for our studies.

**FIG 1.**
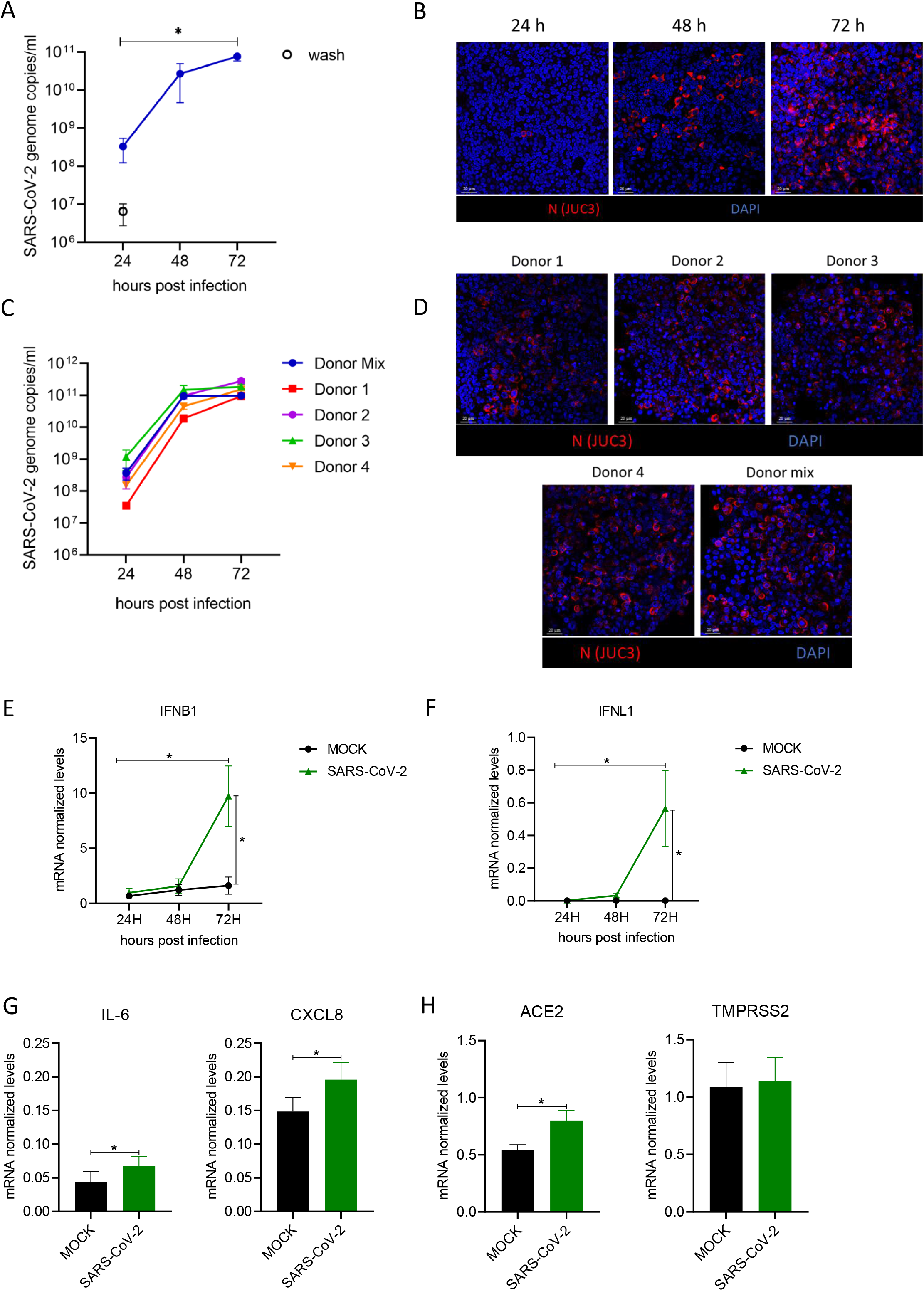
Characterization of a SARS-CoV-2 infection model using air-liquid interface cultures of primary bronchial epithelial cells. Mixes of PBEC derived from 4-5 individual donors were cultured for 5 weeks at ALI before they were infected with SARS-CoV-2 (30,000 PFU per insert). (A) The viral load at 24, 48 and 72 hpi in cultures was determined by quantifying the number of extracellular viral RNA copies by RT-qPCR. The open circle represents the amount of (input) viral RNA that remained after washing the inserts at 2 hpi. Data are mean ± SEM. n=4 independent experiments with the same donor mix. (B) For immunofluorescence microscopy, cells were fixed at 72 hpi and (double) labelled with rabbit polyclonal anti-SARS-CoV-2 N protein antibody (JUC3) and with 4’,6-diamidino-2-phenylindole (DAPI) for nuclear staining. Images shown at 400 x original magnification are representative for results obtained with cells from 3 independent experiments. (C-D) The viral load in cultures derived from single donors or the mix of these donors (blue) was determined by quantifying the number of extracellular viral RNA copies by RT-qPCR. The infected cells in the cultures were visualized by immunofluorescence microscopy at 72 hpi. (E-H) Analysis of gene expression of *IFNB1* (E), *IFNL1* (F), *IL-6* and *CXCL8* (IL-8) (G), *ACE2* and *TMPRSS2* (H) normalized to two reference genes (*RPL13A/ATP5B*) in mixes of 5 donors by RT-qPCR. The graphs represent the mRNA levels at 72 hpi. Data are mean values ± SEM. n=3 independent experiments derived from the same donor mix cultured for at least 4 weeks in E and F, and n=5 independent experiments derived from the same donor mix and 2 individual donors cultured for at least 4 weeks in G and H. Statistical analysis was conducted using two-way ANOVA with a Tukey/Bonferroni post-hoc test. Significant differences are indicated by *P<0.05.

To validate the relevance of our cell culture model for studying the epithelial defense against SARS-CoV-2 infection, we measured antiviral responses (IFN-β1 and IFN-λ1), inflammatory cytokines [IL-6 and IL-8(CXCL8)] as well as expression of SARS-CoV-2-entry factors (ACE2 and TMPRSS2) in the donor mix cultures. We observed that SARS-CoV-2 infection did not affect mRNA levels of *IFN-B1* and *IFN-L1* at 24 and 48 h, but strongly increased their expression at 72 h post-infection (Fig. 1E and 1F). At 72 hpi, mRNA levels of both *IL-6*, *IL-8* and *ACE2* displayed a modest but significant increase in SARS-CoV-2-infected cultures (Fig. 1G, H), while the expression of *TMPRSS2* remained unchanged (Fig. 1H). These findings show reproducible and robust infection kinetics in our model, as well as a significant but late epithelial antiviral and inflammatory response that is characteristic in SARS-CoV-2 infection (25). Using ALI-PBEC cultures of mixed donor cells proved to be a practical way to investigate SARS-CoV-2 infection of airway epithelial cell cultures in a model that incorporates genetic variation between individuals, while limiting sample size to allow investigation of many variables.

### Effect of culture duration of primary bronchial and tracheal epithelial cell cultures on SARS-CoV-2 infection

We compared infection of ALI-PBEC and ALI-PTEC cultures as these epithelial tissues differ in cellular composition as a result of their disparate anatomical origin. To investigate if duration of the ALI culture has an effect on the infection kinetics of SARS-CoV-2, we allowed ALI-PBEC and ALI-PTEC to differentiate for 3, 4 or 5 weeks, after which they were infected with 30,000 PFU of SARS-CoV-2. Viral load was analyzed at 72 hpi (Fig. 2). In both ALI-PBEC and ALI-PTEC, an increase in intra- and extracellular viral RNA as well as infectious virus particles was observed with longer culture time, with the highest viral load observed at 5 weeks after ALI (Fig. 2A-2C). A gradual 1-2 log increase in SARS-CoV-2 progeny production was observed from 3 to 5 weeks of differentiation when three independent cultures of different PTEC and PBEC donor mixes were followed over time (Fig. S1). This shows the robustness of our model and the reproducibility of our results, independent of the ALI-PBEC or ALI-PTEC donor mix used. Immunofluorescence staining of these cultures for the viral nucleocapsid protein also demonstrated an increase in the number of infected cells with increasing culture duration (Fig. 2D). A significantly higher viral load was observed in ALI-PBEC than in ALI-PTEC, with an average 10-fold difference in extracellular SARS-CoV-2 RNA copies (Fig. 2A) and infectious progeny (Fig. 2B), in particular in 5-week differentiated cultures.

**FIG 2.**
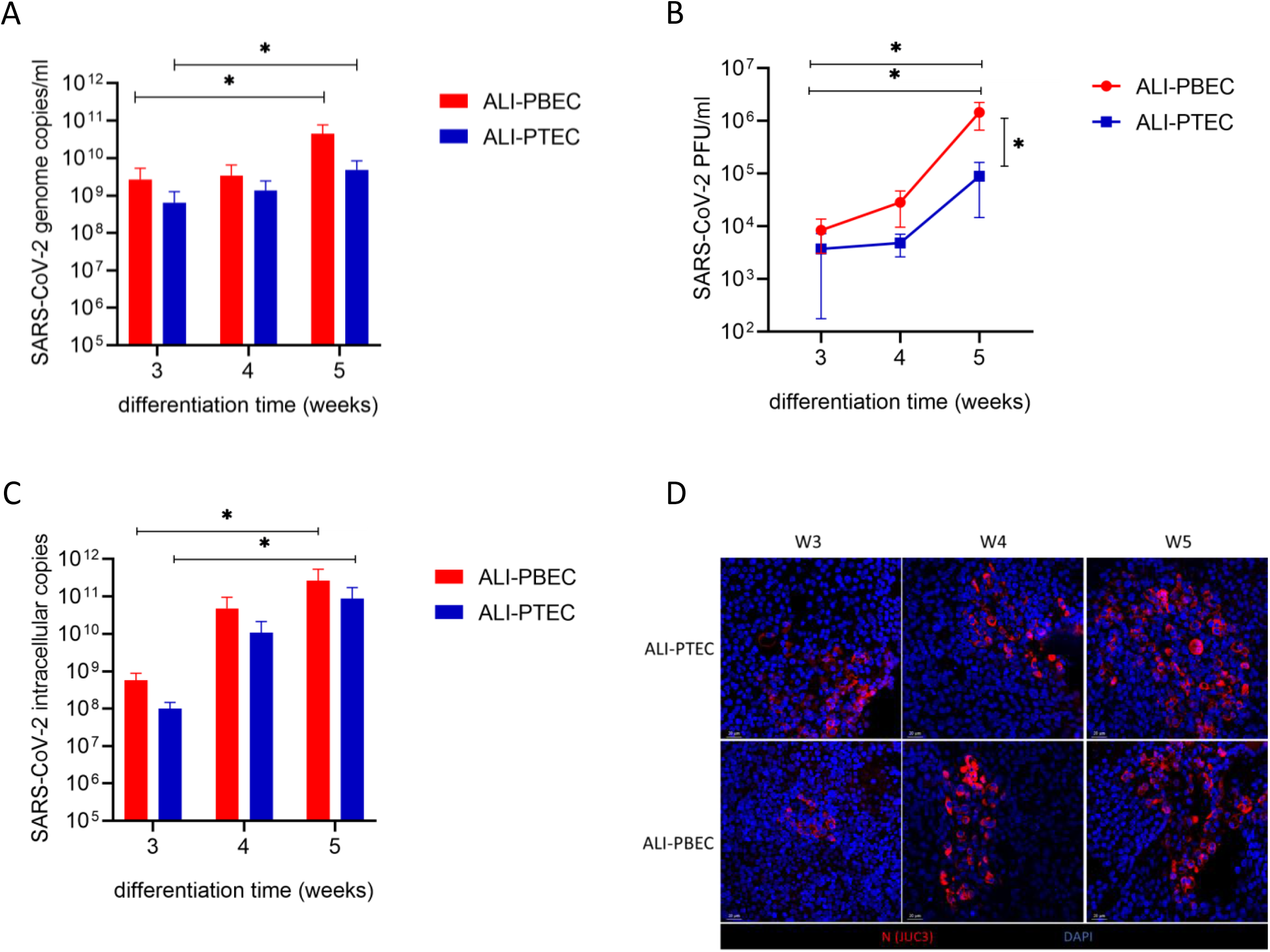
Effects of culture duration on SARS-CoV-2 infection in PTEC and PBEC. ALI-PBEC/ALI-PTEC (mix of 5 donors) cultured for 3-5 weeks were infected with SARS-CoV-2 (30,000 PFU per insert). (A) Extracellular viral RNA copies in the apical wash or (C) intracellular copies were measured by RT-qPCR. (B) Viral infectious progeny was determined by plaque assay in Vero E6 cells. Mean values ± SEM is presented from 3 independent experiments using 3 different donor mixes. Statistical analysis was conducted using two-way ANOVA with a Tukey/Bonferroni post-hoc test. Significant differences are indicated by *P<0.05. (D) Cells were immunofluorescence stained with rabbit polyclonal anti-SARS-CoV-2 N protein antibody (JUC3) and DAPI for nuclear staining. Images shown are representative for results obtained with ALI-PBEC and ALI-PTEC from the same 3 independent experiments shown in A-C at 400 x original magnification.

In conclusion, our results show that the anatomical origin of the epithelial cells and the culture duration have an effect on the levels of infection, suggesting that cellular composition is a contributing factor. To further investigate this possibility, we identified the cell types that were primarily infected by SARS-CoV-2 and assessed their abundance with respect to culture duration.

### SARS-CoV-2 primarily infects ciliated and goblet cells in primary human bronchial and tracheal epithelial cell cultures

Previous studies have indicated that within the human respiratory tract, predominantly ciliated cells, but also goblet cells of the airway epithelium can be infected with SARS-CoV-2 (14), besides alveolar epithelial cells (26, 27). To assess if ciliated and goblet cells were also the target cells in our cultures, we investigated the colocalization of the SARS-CoV-2 nucleocapsid protein with either acetylated α-tubulin as a marker for ciliated cells, or MUC5AC as a marker for goblet cells.

We observed positive staining of the viral nucleocapsid protein in acetylated α-tubulin-positive ciliated cells in ALI-PBEC (Fig. 3A) and also in MUC5AC-positive goblet cells (Fig. 3B), suggesting that both cell types are indeed infected by SARS-CoV-2. Similar results were obtained with ALI-PTEC (Fig. S2), indicating that goblet cells and ciliated cells originating from both anatomical locations in the airways can be infected.

**FIG 3.**
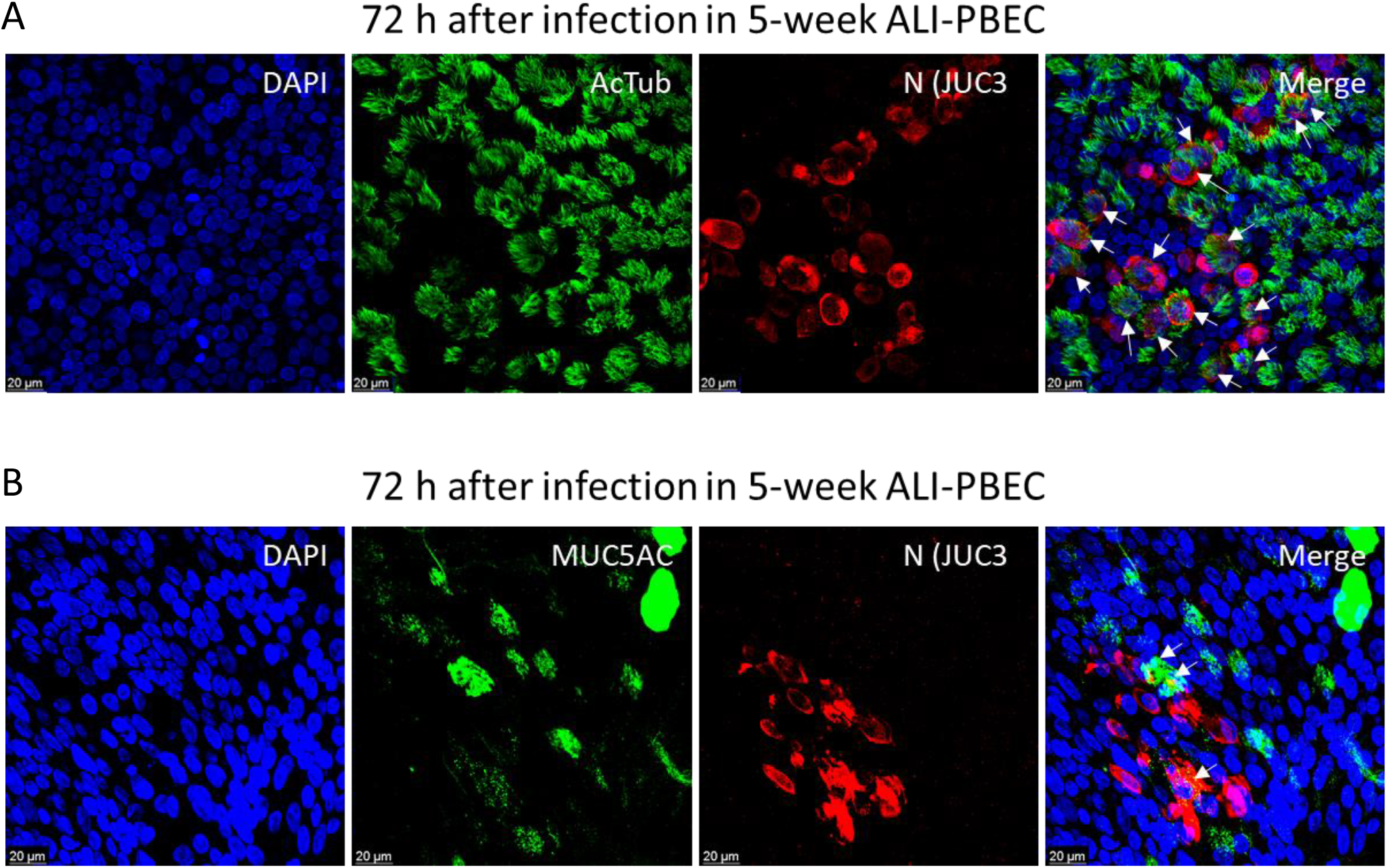
Cell types infected by SARS-CoV-2 in ALI-PBEC. ALI-PBEC (mix of 5 donors) cultured for 5 weeks were infected with SARS-CoV-2 (30,000 PFU per insert). Immunofluorescence staining at 72 hpi with primary antibodies against acetylated α-tubulin as a ciliated cell marker (A) or MUC5AC as a goblet cell marker (B) in combination with anti-SARS-CoV-2 N protein antibody (JUC3) and DAPI for nuclear staining. Immunofluorescence images shown are representative for results of 3 independent experiments derived from the same donor mix with 630 x original magnification. White arrows indicate cells that display both green and red labelling.

### Cellular composition of primary human airway epithelial tissues differs depending on time of culturing

Since we found that increased viral infection with prolonged culture time and observed that SARS-CoV-2 targeted both ciliated and goblet cells, we hypothesized that changes in cellular composition might account for these differences. Therefore, we compared cellular composition between 2, 3, 4 and 5 week-differentiated cultures, for the same donor mixes that were used for infection. The results showed that the cultures were well-differentiated at all time points, showing expression of markers related to all cell types (ciliated, goblet, club and basal cells) in ALI-PBEC and PTEC (Fig. 4 and S3). However, there were differences in the proportions of goblet and ciliated cells. Using FOXJ1 and acetylated α-tubulin as markers for ciliated cells, we observed that the percentage of FOXJ1-positive cells was significantly higher in ALI-PBEC after 5 weeks ALI culture compared to 3-week cultures. The percentage of ciliated cells was significantly higher in ALI-PBEC than in ALI-PTEC at all culture durations (Fig. 4A and 4B). The change in the percentage of MUC5AC-positive goblet cells was not significant over time in ALI-PBEC and ALI-PTEC (Fig. 4A and 4B). The sum of the percentage of ciliated and goblet cells was higher in week 5 ALI-PBEC cultures compared to week 3 cultures, and it was also higher in ALI-PBEC than ALI-PTEC when 5-week cultures were compared (Fig. 4B). These differences in the total proportion of both cell types between PTEC and PBEC and between cultures of different duration correlated with differences in virus levels (Fig. 2). Furthermore, mRNA levels of *FOXJ1* were significantly increased in 4 week ALI-PBEC compared to 3 week cultures, however they did not further increase in 5 week cultures (Fig. 4C). In addition, *FOXJ1* mRNA was higher in 4/5-week ALI-PBEC cultures compared to 4/5-week PTEC cultures (Fig. 4C). In line with these findings, *MUC5AC* mRNA levels were higher at week 5 in ALI-PTEC cultures compared to week 3, and also higher than in week 5 ALI-PBEC (Fig. 4C). In contrast, there was no significant difference in the expression of *SCGB1A1* (club cell marker) and *TP63* (basal cell marker) (Fig. S3). These results suggest that despite the early presence of transcripts which are specific for certain cell types, maturation of these cell types (which also requires expression at the protein level) continues for several weeks in cultures at ALI. In addition to culture duration, difference of ciliated and goblet cell markers between ALI-PBEC and ALI-PTEC indicated that the origin of the epithelial cells (tracheal versus bronchial) also has an impact on epithelial cellular composition. Epithelial cellular composition, and especially percentages of ciliated and goblet cells were associated with increased SARS-CoV-2 infection. While the observed increase in viral load cannot be attributed to one cell type alone, the levels of both ciliated and goblet cells appear to affect infection outcome in our model.

**FIG 4.**
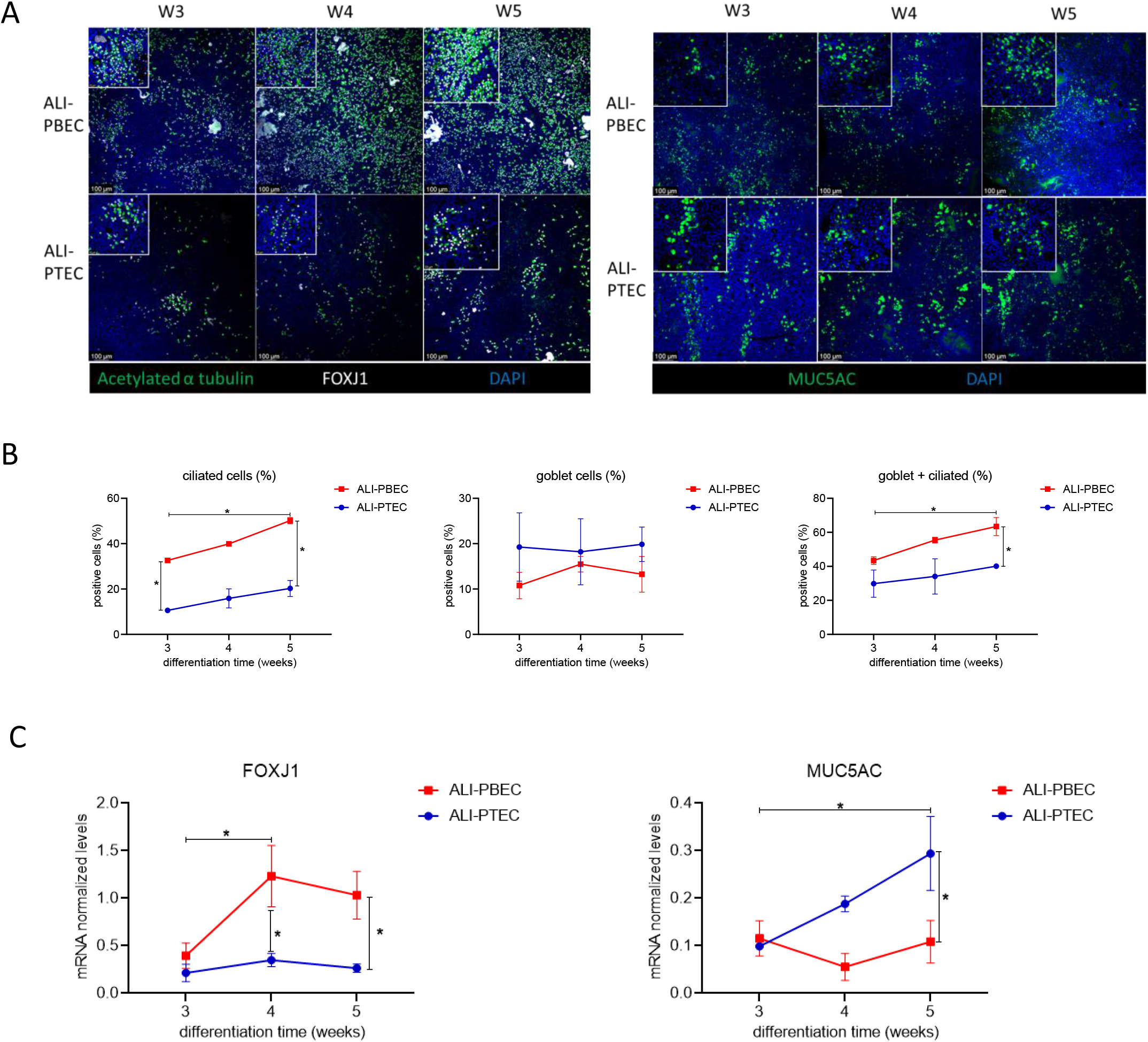
Effect of culture duration on epithelial differentiation markers in PTEC and PBEC. ALI-PTEC/PBEC (mix of 5 donors) were differentiated at ALI for 3, 4 or 5 weeks, fixed and analyzed by immunofluorescence microscopy (A) using antibodies against acetylated α-tubulin and FOXJ1 (ciliated cell markers) or MUC5AC (goblet cell marker) in combination with DAPI for nuclear staining. Images shown are representative for results of 3 independent experiments with 100x/400x (insert) original magnification. (B) Quantification of FOXJ-positive cells and MUC5AC-positive cells was done by Image J software. (C) mRNA levels of *FOXJ1* and *MUC5AC* were measured by RT-qPCR. n=3 independent experiments derived from 3 different donor mixes same as in FIG 2. Data are mean ± SEM. Analysis of differences was conducted using two-way ANOVA with a Tukey/Bonferroni post-hoc test. Significant differences are indicated by *P<0.05.

### Modulating epithelial cellular composition has moderate effects on SARS-CoV-2 infection

To further explore the association between epithelial cellular composition and viral infection, we skewed differentiation of ALI-PBEC during the last 2 weeks of culturing either towards an enrichment in ciliated cells, using the γ-secretase inhibitor DAPT (that acts as an inhibitor of Notch signaling), or towards an enrichment in goblet cells using interleukin 13 (IL-13) (22–24). We infected these cultures with SARS-CoV-2 to analyze if enrichment of one of the cell types impacted infection kinetics. We could verify that DAPT treatment caused a marked increase in ciliated cells (FOXJ1 and acetylated α-tubulin positive) while the number of goblet cells (MUC5AC positive) was decreased (Fig. 5A and S4A). Conversely, IL-13 treatment increased the fraction of goblet cells and decreased the number of ciliated cells (Fig. 5A and S4A). This was confirmed by quantifying expression levels of mRNAs that serve as markers for each cell type (Fig. S4B). Although DAPT and IL-13 treatment caused a shift in the ratio between ciliated and goblet cells, it did not change the combined/total number of these two cell types (Fig. S4A). Notably, there were almost no goblet cells present in DAPT-treated cultures.

**FIG 5.**
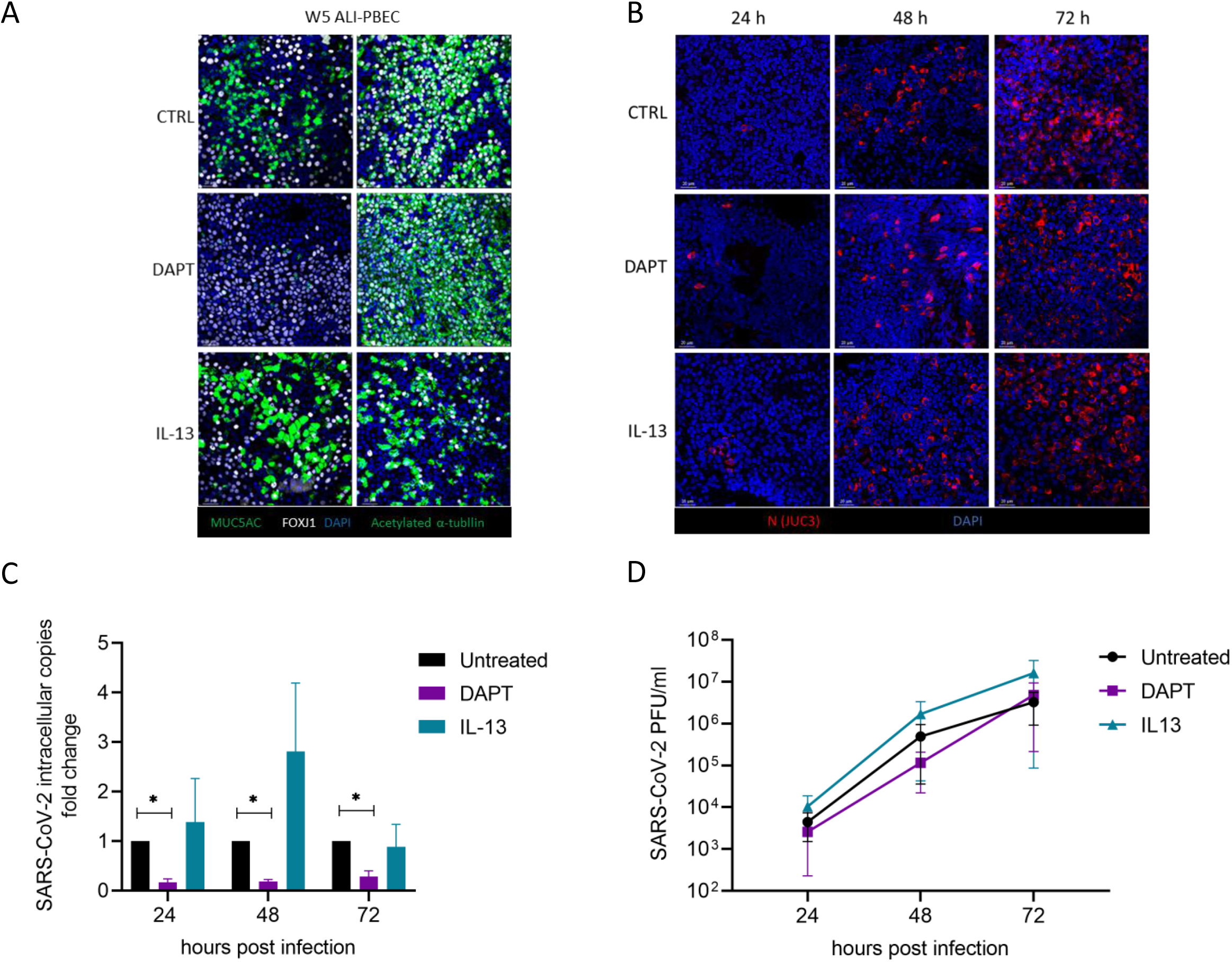
Effect of IL-13 treatment, and DAPT-mediated inhibition of Notch signaling on epithelial susceptibility to SARS-CoV-2 infection. ALI-PBEC (mix of 4-5 donors) were differentiated for 3 weeks, before addition of DAPT (5 µM) or IL-13 (1 ng/ml) and differentiation for an additional 2 weeks. (A) After in total 5 weeks of differentiation, ALI-PBEC were fixed, stained using primary antibodies against MUC5AC and FOXJ1 (goblet cell marker, ciliated cell marker) or acetylated α-tubulin together with FOXJ1 (ciliated cell markers) in combination with DAPI for nuclear staining and analyzed by immunofluorescence microscopy. (B) SARS-CoV-2 infected cells were stained with primary antibodies against SARS-CoV-2 N protein (JUC3) in combination with DAPI for nuclear staining. (C) Intracellular viral RNA copies were measured by RT-qPCR and their fold change in treated cultures compared to untreated controls was determined. (D) Plaque assay was performed to titrate viral progeny in the apical washes. n=3 independent experiments derived from 3 different donor mixes. Data are mean ± SEM. Analysis of differences was conducted using two-way ANOVA with a Tukey/Bonferroni post-hoc test. Significant differences are indicated by *P<0.05 compared with untreated samples.

We next infected the DAPT- or IL-13-treated cultures with SARS-CoV-2 and infection kinetics were analyzed. In line with the data shown in Fig. 1, only a few infected cells were identified at 24 hpi in untreated and treated cultures by immunofluorescence staining (Fig. 5B). At 48 hpi, DAPT-treated cell cultures showed a lower number of infected cells compared to untreated or IL-13-treated cells (Fig. 5B). In untreated, DAPT- and IL-13-treated cultures, spread of infection was observed and a substantial fraction of the cells was infected at 72 hpi, with IL-13-treated cell cultures displaying the highest level of infected cells (Fig. 5B). The level of intracellular SARS-CoV-2 RNA was lower in DAPT-treated cells at all time points compared to the controls (Fig. 5C). We did not observe a significant decrease of infectious viral particles in DAPT-treated cultures (Fig. 5D). These findings suggested that a DAPT-induced increase in ciliated cells did not enhance infection, but rather was associated with decreased viral replication or spread. Treatment with IL-13 on the other hand showed a trend towards an increase in intracellular copies and infectious virus particles compared to untreated and DAPT-treated cultures (Fig. 5C and 5D).

To exclude the possibility that the effects observed in DAPT-treated cultures were a direct consequence of inhibition of Notch signaling rather than epithelial remodeling, we treated cells with DAPT either starting 24 h before (and during) infection (a time period considered insufficient to cause a shift in epithelial differentiation) or directly after infection. These ‘short term treatments’ with DAPT did not result in significant changes in the amount of intracellular viral RNA copies or extracellular viral load, suggesting that inhibition of Notch signaling has no direct effect on SARS-CoV-2 replication (Fig. S4C). These findings indicate that the effects of DAPT treatment on infection are more likely due to an effect on cellular composition.

### Culture duration does not affect SARS-CoV-2-entry factors

Viral entry factors are expressed by human airway epithelial cells and their expression varies between different airway epithelial cell types (18). Since cells could be infected more efficiently when they were allowed to differentiate for longer periods of time, we investigated if this could (also) be related to changes in expression of receptors used by SARS-CoV-2 to enter the cells. SARS-CoV-2 uses mainly ACE2 as its receptor, but studies demonstrated that other co-factors can also determine its cellular tropism (19). We found that culture duration did not affect gene expression of *ACE2*, *TMPRSS2*, *CD147* and *GRP78* (Fig. 6A) These findings indicated that differences in susceptibility to SARS-CoV-2 infection between ALI-PBEC and ALI-PTEC, and between 3-, 4- and 5-week differentiated cultures were likely not due to differences in epithelial expression of SARS-CoV-2 cell entry-related genes. Furthermore, also treatment with DAPT or IL-13 did not markedly affect expression of viral entry factors (Fig. 6B).

**FIG 6.**
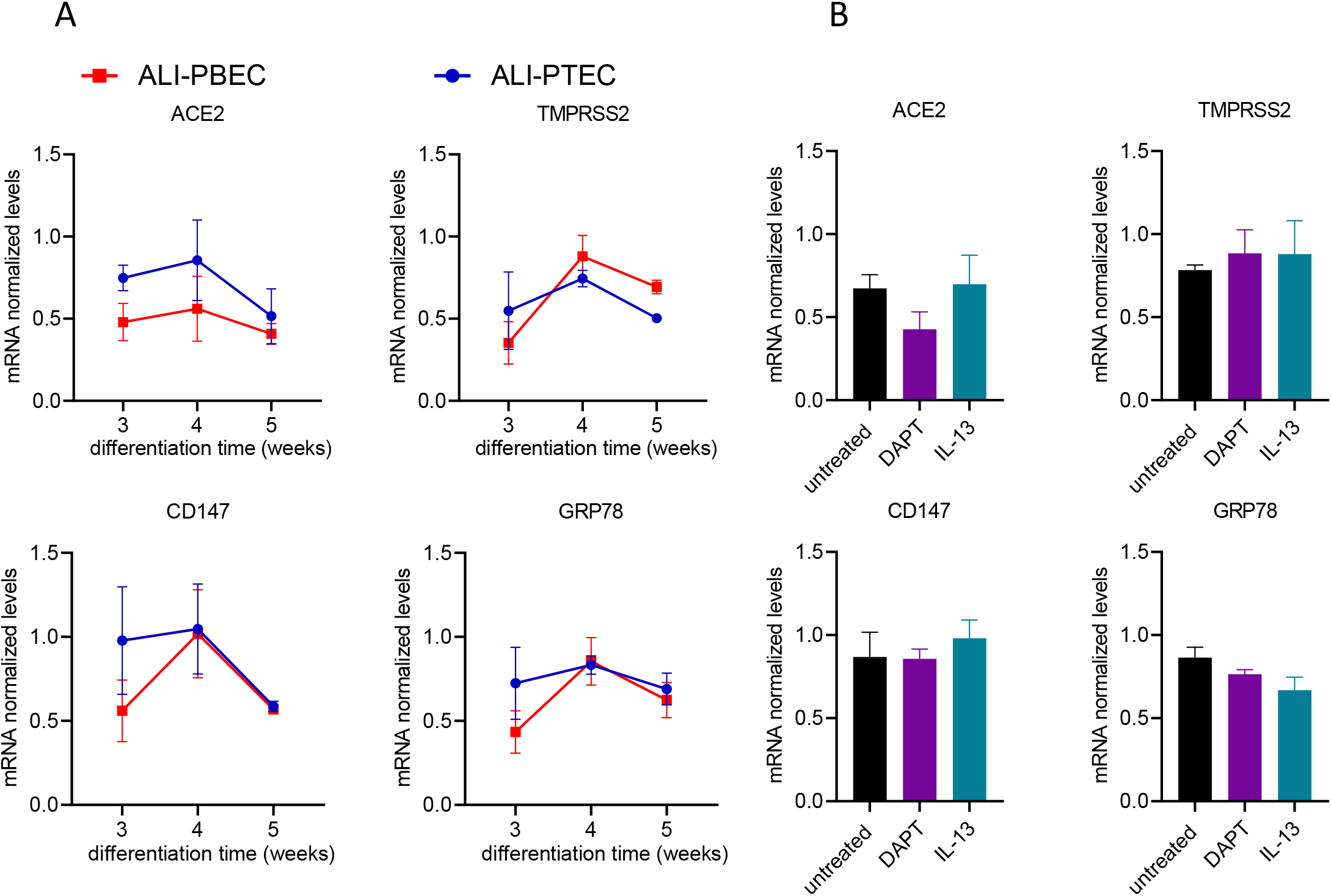
Effect of culture duration on expression of SARS-CoV-2 cell-entry related factors in well-differentiated primary epithelial cells. ALI-PBEC/PTEC cultured for 3-5 weeks with or without the presence of DAPT/IL-13 (during the last 2 weeks) were infected with SARS-CoV-2 (30,000 PFU per insert). Expression of *ACE2*, *TMPRSS2*, *CH147* and *GRP78* over time (A) or in DAPT/IL-13 treated cultures (B) were measured by RT-qPCR. n=3 independent experiments using 3 different donor mixes. Data are mean values ± SEM. Analysis of differences was conducted using two-way ANOVA with a Tukey/Bonferroni post-hoc test or paired t test.

### SARS-CoV-2-induced antiviral responses in primary human airway epithelial cultures

Antiviral responses in the epithelium are critical for protection against viral infections. Therefore, we investigated whether there were differences in antiviral responses depending on the epithelial culture duration, which might explain the observed increased susceptibility to infection with longer culture times. In ALI-PTEC, SARS-CoV-2-induced mRNA levels of both *IFNB1* and *IFNL1* increased significantly with culture duration. When cultures were infected at week 3 or 4 there was little *IFNB1* or *IFNL1* mRNA produced, while expression of these genes was strongly upregulated by SARS-CoV-2 infection in 5 week old cultures (Fig. 7A and 7B). Gene expression analysis of *IFNB1* and *IFNL1* showed a similar increasing trend in ALI-PBEC upon infection (Fig. 7C and 7D). When 5-week PBEC were (long-term) treated with DAPT, we observed lower SARS-CoV-2-induced antiviral responses than in untreated and IL-13-treated cultures (Fig. 7E and 7F). All these findings correlate with the observed differences in the number of infected cells and viral load resulting from culture duration and DAPT and IL-13 treatment (Fig. 5). This suggests that antiviral responses were likely not affected by differentiation time or long-term DAPT/IL-13 treatment, but rather by a direct response to the virus load.

**FIG 7.**
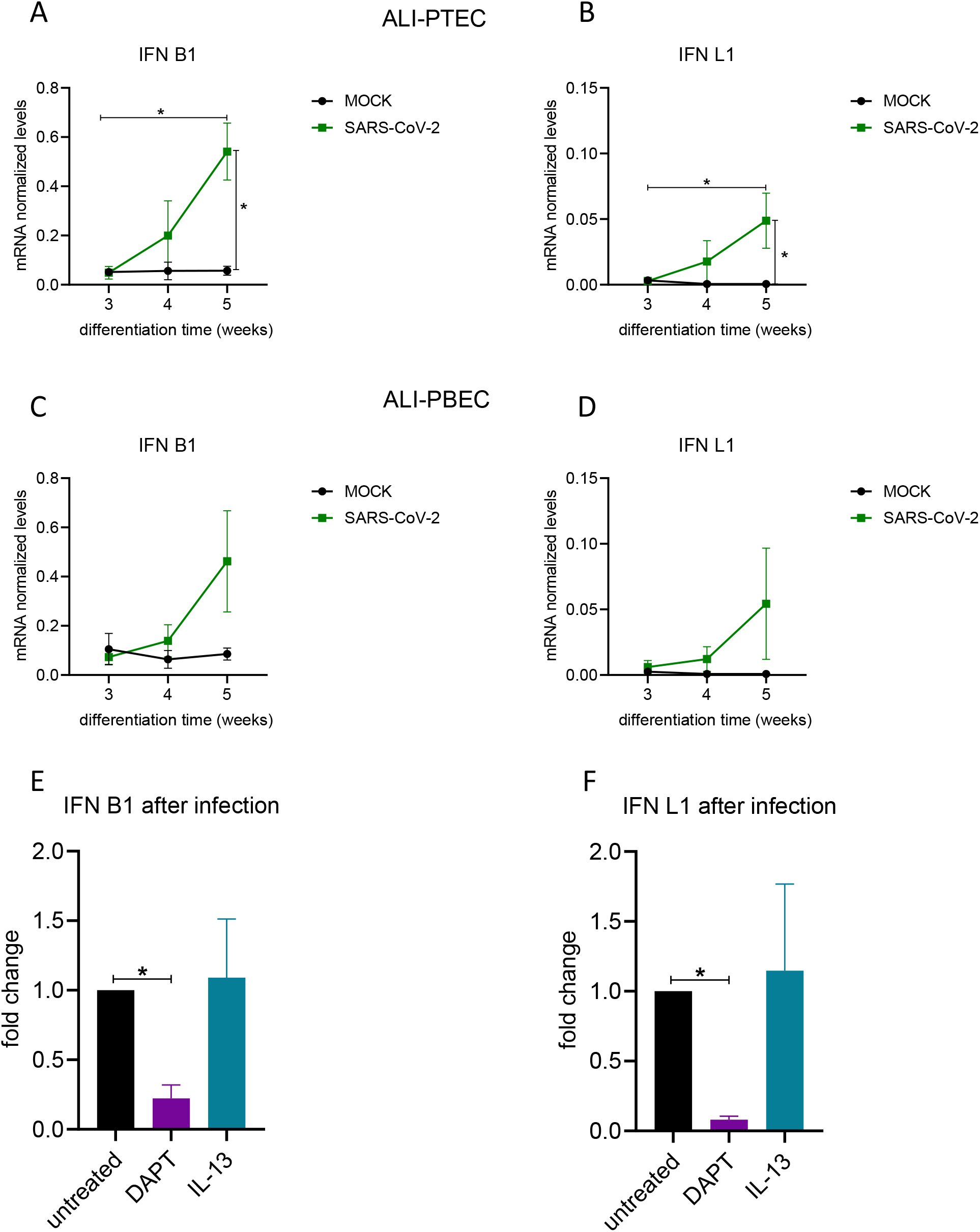
Antiviral responses after SARS-CoV-2 infection. ALI-PTEC/PBEC cultured for 3-5 weeks (A-D), and 5-week ALI-PBEC (E, F) cultured at ALI with or without the presence of DAPT/IL-13 (during the last 2 weeks), were infected by SARS-CoV-2 (30,000 PFU per insert). Cells were lysed at 72 hpi to quantify mRNA levels of *IFNB1* (A/C/E) and *IFNL1* (B/D/F) by RT-qPCR. n=3 independent experiments using 3 different donor mixes. Data are mean ± SEM. Fold change in E and F was compared to untreated controls. Analysis of differences was conducted using two-way ANOVA with a Tukey/Bonferroni post-hoc test. Significant differences are indicated by *P<0.05.

## DISCUSSION

Here we describe and characterize the use of well-differentiated human primary airway epithelial cell cultures to investigate the effect of cell composition on susceptibility to SARS-CoV-2 infection. Our key finding is that culture duration at the air-liquid interface, which is needed to achieve mucociliary differentiation, is an important contributor to SARS-CoV-2 infection kinetics. Specifically, the percentage of goblet and ciliated cells is pivotal as these cell types are likely the first to become infected.

As a highly relevant cell model, cultures of well-differentiated airway epithelial cells are employed to study infection biology of SARS-CoV-2. With regard to differentiation time, there is no consensus or standardized protocol for ALI-PBEC cultures and it is described in literature as anywhere from 2 to 6 weeks after start of culture at the air-liquid interface (14, 15, 28). To validate our model, we decided to use up to 5 weeks differentiated ALI-PBEC cultures, in line with a recent study (14). In this study, we report efficient infection with the peak of viral load at 72 hpi with around 10^11^ extracellular RNA copies and 10^6^ PFU/ml, and observed virus spread across the epithelium by immunofluorescence staining of infected cells. The amount of viral progeny produced is in line with other studies assessing viral load levels (14). We have established our findings using cultures derived from single donors and mixes derived from multiple donors, as well as compared primary bronchial and tracheal cell cultures. The viral load was previously reported to plateau at 72 hpi and onset of related antiviral responses as well as inflammation were also delayed, and therefore we mainly focused on this time point for analyses (14, 15). In this study, mRNA levels of *IL-6* and *IL-8* were significantly upregulated by SARS-CoV-2, which is consistent with previous findings (29) and different from results found in human bronchial epithelial cell line (16HBE) (30). These findings highlight the importance of using well-differentiated cell models for SARS-CoV-2 infection.

Multiple variables can contribute to epithelial susceptibility to SARS-CoV-2 infection, and here we hypothesized cellular composition as being one of them. Epithelial cellular composition is inherently different throughout the human respiratory tract (10, 11). Ciliated, goblet, club and basal cells are the main cell types of the airway epithelium, which vary upon culturing due to different stages of differentiation (31). To elucidate the impact of cellular composition on infection kinetics, we used ALI-PTEC and ALI-PBEC cultured for 3, 4 or 5 weeks. Firstly, we measured higher viral loads in ALI-PBEC than in ALI-PTEC. Furthermore, we found increased infection with prolonged culture duration in PBEC and PTEC.

To investigate the role of cellular composition in these observations, we first assessed which cells were infected by SARS-CoV-2. Recent studies looking into the cell tropism of SARS-CoV-2 have shown infection of multiple epithelial cell types, among them ciliated cells, goblet cells and club cells of the airway epithelium and type 2 alveolar epithelial cells (14, 15, 32, 33). In line with other studies, we found that both ciliated and goblet cells were infected by SARS-CoV-2 in our ALI-PBEC and ALI-PTEC cultures, suggesting both ciliated cells and goblet cells support viral replication.

We next compared cellular composition in ALI-PBEC and ALI-PTEC cultured for 3-5 weeks, and found that differences in ciliated and goblet cell numbers did not explain the differences in infection. Our data suggest that the total percentages of target cells that might be initially susceptible, i.e. ciliated and goblet cells, influences the efficiency of infection. Interestingly, we also observed cells co-expressing markers of ciliated cells (FOXJ1) and goblet cells (MUC5AC) in our culture. This specific cell population, which was reported by e.g. Garcia *et al.* (31) and is suggested to represent a transitional state between goblet cells and ciliated cells, was recently labeled as transient secretory cells (34). Based on their relative high expression of ACE2 and TMPRSS2, Lukassen et al(34) suggested that these transient secretory cells may be particularly vulnerable to SARS-CoV-2 infection.

Unexpectedly, skewing differentiation towards ciliated cells by treatment of our cultures with DAPT, actually decreased infection compared to untreated and IL-13-treated cultures, highlighting the importance of the presence of goblet cells, besides ciliated cells. Of note, there was a very weak signal of the goblet cell marker (MUC5AC) after treatment with DAPT, and therefore, the presence of some goblet cells in these cultures cannot be excluded. Another explanation would be that ciliated cells catch the viruses through cilia, and mucus produced by goblet cells helps spread the infection after release and therefore hinders clearance by ciliated cells, which would be consistent with previous findings that SARS-CoV-2 infects mucus-covered ciliated cells (35). To exclude a possible direct effect of DAPT treatment on viral replication via inhibition of Notch signaling (36) independent of formation of ciliated cells, short-term treatment with DAPT during infection was performed. However, such acute treatment did not alter viral infection.

Conversely, (slightly) increased viral loads were observed in IL-13-treated cultures compared to control and DAPT-treated cultures, in line with the suggestion that patients with allergic asthma, a disease associated with IL-13-induced changes in epithelial cells, may be somewhat more susceptible to SARS-CoV-2 infection (37). Collectively, these observations indicate that DAPT- and IL-13-mediated modulation of epithelial cell differentiation does not provide a simple answer to linking differences in epithelial cell composition to SARS-CoV-2 infectivity, but hints at the role of other factors like mucus secretion of goblet cells (38), which could hinder clearance of the virus in the epithelium, or ciliary (dys)functioning that may influence spread of the virus (39).

We excluded the possibility that the abundance of viral entry factors contributed to the observed differences in SARS-CoV-2 replication/spread, as their expression was not significantly higher in ALI-PBEC than ALI-PTEC and did not increase with the time of differentiation. Zhang *et al.* showed that gene expression of *ACE2* was increased in differentiated ALI cultures compared to submerged cultured basal cells and that their levels did not change during differentiation (28), which is in line with our findings. Furthermore, DAPT treatment and IL-13 treatment did not change the levels of viral entry factors in our study. However, recent studies showed that long-term IL-13 exposure decreased, while DAPT increased ACE2-positive apical cells in nasal epithelial cell cultures, differentiated for 3 weeks (40). Differences with our findings could be explained by a possible discrepancy between mRNA and protein levels or differences in cell culture duration.

Another factor that shapes infection is the antiviral response. Recent studies showed that SARS-CoV-2 upregulated type III IFN (IFN-λ1) in human intestinal epithelial cells and an organoid-derived bronchioalveolar model (relatively late in infection) (25, 41). Importantly, addition of type III IFN-λ1 inhibited SARS-CoV-2 infection in human Calu-3 and simian Vero E6 cells in a dose dependent manner (42). In this study, we found that expression of type I and III IFNs showed a similar pattern as viral replication over culture time in ALI cultures, which indicated that antiviral responses may not contribute to the gradual increase in viral infection with prolonged culture time.

Our comparison of mixed donor cultures to cultures of single donors showed similar infection kinetics, which is in line with our previous findings (12). This demonstrates that the use of mixed donor cultures is an efficient way to recapitulate natural variability between donors while saving on resources (cells, culture plastics and media) that are in high demand/short supply. The level of donor-to-donor variation was also represented in inter-experimental variation when using the same donor mix. However, there are also some limitations in this study, like the sources of cells. PBEC were derived from tumor-free resected bronchial tissue from (ex-)smoking non-COPD patients with lung cancer, and PTEC were from lung donors without lung disease, which may have affected the comparison between PBEC and PTEC.

In summary, we show that culture time of airway epithelial cells at the air-liquid interface is a determinant of their susceptibility to SARS-CoV-2 infection, which can only be partly explained by differentiation status based on the amount of ciliated and goblet cells. Differences in expression of entry-associated factors, like ACE2 or TMPRSS2 do not explain the increased susceptibility of airway epithelial cell cultures upon prolonged culture. Our observation that IL-13 treatment causes a limited increase in SARS-CoV-2 infection may be relevant for understanding the impact of type 2 allergic inflammation on SARS-CoV-2 susceptibility. Decreased infection following prolonged inhibition of Notch signaling by DAPT highlights the importance of the presence of both ciliated and goblet cells and warrants further investigation. Ultimately, cellular maturation/differentiation seems intertwined with virus load and spread of the infection over the culture. The proposed cell culture set-up provides a robust tool to test antiviral compounds and acquire additional insight into infection biology of SARS-CoV-2.

## MATERIALS AND METHODS

### Cell culture

PBEC were isolated from tumor-free resected bronchial tissue that was obtained from patients undergoing resection surgery for lung cancer at the Leiden University Medical Center (Leiden, the Netherlands). Use of such lung tissue that became available for research within the framework of patient care was in line with the “Human Tissue and Medical Research: Code of conduct for responsible use” (2011) (www.federa.org), that describes the opt-out system for coded anonymous further use of such tissue. PTEC were isolated from residual tracheal and main stem bronchial tissue from lung transplant donors post-mortem at the University Medical Center Essen (Essen, Germany). Use of such donor tissue for research was approved by the ethical committee of the Medical faculty of the University Duisburg-Essen (ID: 19-8717-BO). After isolation, cells were expanded in T75 flasks and frozen in liquid nitrogen until use as previously described (43).

To achieve mucociliary differentiation, primary human bronchial/tracheal epithelial cells were cultured at the air-liquid interface (ALI) as previously described (22). Briefly, approximately 150,000 cells (30,000 cells/donor when mixing cells from 5 donors and 40,000 cells/donor when using 4 donors) were seeded onto 12-insert Transwell membranes (Corning Costar, Cambridge, MA, USA), which were coated with PBS supplemented with 5 μg/ml human fibronectin (Promocell, Heidelberg, Germany), 30 μg/ml PureCol (Advanced BioMatrix, CA, USA) and 10 μg/ml bovine serum albumin (Fraction V; Thermo Fisher Scientific, Carlsbad, CA, USA), in a 1:1 mixture of Bronchial Epithelial Cell Medium-basal (BEpiCM-b; ScienCell, Sanbio) and Dulbecco’s modified Eagle’s medium (DMEM) (Stemcell Technologies, Köln, Germany), further referred to as B/D medium. This B/D medium contains 12.5 mM Hepes, bronchial epithelial cell growth supplement, 100 U/ml penicillin, 100 ug/ml streptomycin (all from ScienCell), 2 mM glutaMAX (Thermo Fisher Scientific). B/D medium was supplemented during submerged culture with 1 nM EC23 (light-stable retinoic acid receptor agonist; Tocris, Abingdon, UK). After confluence was reached, the apical medium was removed and cells were cultured at the ALI in B/D medium with 50 nM EC23 for 3-5 weeks, medium was refreshed and the apical side was washed three times a week with warm PBS to remove excess mucus.

PBEC (40,000 cells) from the individual donors were expanded in separate cultures, before mixing all donors when seeding them on inserts. For these donor mixes, higher cell numbers for plating on inserts were used when compared to using individual donors to avoid selective advantage of possible faster-proliferating cells of specific donors. To shift cell differentiation towards an increased number of goblet or ciliated cells, ALI-PBEC were incubated in BD medium supplemented with 50 nM EC23, and either 1 ng/ml IL-13 (Peprotech) or 5 µM DAPT (γ-secretase inhibitor, TOCRIS) from day 22 to day 35 culture time.

For acute treatment of DAPT, ALI-PBEC after 5 week culture were either pre-treated with DAPT (5 µM) for 24 hpi or DAPT was directly added (5 µM) after infection.

Vero E6 cells (master stock MM-3 from dept of Medical Microbiology collection, characterized by full genome sequencing) were maintained in Dulbecco’s modified Eagle’s medium with 4.5 g/l glucose with L-Glutamin (DMEM; Lonza), supplemented with 8% fetal calf serum (FBS; CapriCorn Scientific) and 100 U/ml of penicillin/streptomycin (Sigma-Aldrich). All cell cultures were maintained at 37°C. Infections for plaque assay in Vero E6 cells were performed in Eagle’s minimal essential medium with 25 mM HEPES (EMEM; Lonza) supplemented with 2% FCS, 2 mM L-glutamine (Sigma-Aldrich), and 100 U/ml of penicillin/streptomycin (Sigma-Aldrich).

### SARS-CoV-2 virus

The clinical isolate SARS-CoV-2/Leiden-0002 was isolated from a nasopharyngeal sample collected at the LUMC (GenBank accession nr. MT510999). Then, SARS-CoV-2/Leiden-0002 was passaged twice in Vero E6 cells to obtain the virus stock used for infection. Virus titers were determined by plaque assay as described before (44). All experiments with infectious SARS-CoV-2 were performed at the Leiden University Medical Center biosafety level 3 facilities.

### SARS-COV-2 infection of ALI-PBEC

Prior to infection, excess of mucus was removed by washing the apical surface of the ALI cultures with 200 µl PBS and aspirating it after a 10-min incubation at 37°C. Basal medium was changed every two days. Cells were infected with 200 µl of inoculum prepared in PBS, containing 30,000 PFU of SARS-CoV-2, per insert for 2 h at 37°C on a rocking platform (estimated MOI of 0.03). PBS was used as a solvent control and in mock-infected cells as inoculum. After removal of the inoculum, the apical side was washed three times with PBS. Then, cells were incubated at 37°C. Viral progeny was harvested from the apical side at 24h, 48h and 72h.p.i. as indicated below. Cells were infected after 3 to 5 weeks of differentiation as indicated. For cells under DAPT or IL-13 treatment, the medium was supplemented with 1 ng/ml IL-13 or 5 µM DAPT after 2 weeks of differentiation, and after 5-week culture time cells were infected. After infection, the basal medium was replaced by fresh B/D medium also supplemented with IL-13 or DAPT.

### RNA isolation, quantitative RT-PCR/real-time PCR (RT-qPCR) and plaque assay analysis

Apical washes were harvested following incubation at 37°C with 200 µl PBS (10 min) and subsequent addition of 800 ul of the TriPure Isolation Reagent (Sigma-Aldrich) to half of apical washes. Tripure reagent was spiked with Equine arteritis virus (EAV) to control for variation in RNA extraction efficiency and possible inhibitors of RT-qPCR. Intracellular RNA was isolated by adding 500 ul of TriPure reagent directly onto the insert. Samples were stored at −20°C until RNA was isolated using the Direct-zol^TM^-96 RNA plate isolation (Zymo), 5PRIME Phase Lock Gel extraction (Quantabio) or Maxwell® 16 simply RNA tissue kit (Promega, the Netherlands). The cellular reference gene PGK-1 was used as a house-keeping gene for intracellular RNA. Primers and probes for EAV and PGK-1 and the normalization procedure was performed as described before (44). Viral RNA was quantified by RT-qPCR using the TaqMan™ Fast Virus 1-Step Master Mix (Thermo Fisher Scientific). Primers and probes were used as described previously (45), but with modifications as listed in Table 1. A standard curve generated by RT-qPCR on 10-fold serial dilutions of a T7 RNA polymerase-generated *in vitro* transcript containing the target sequences was used for absolute quantification of RNA copy numbers. For other gene expression analysis, RNA was reverse-transcribed and cDNA was amplified by real-time qPCR (Bio-Rad, Veenendaal, the Netherlands) using specific primers. Relative normalized gene expression compared to reference genes Ribosomal Protein L13a (RPL13A) and ATP synthase, H^+^ transporting, mitochondrial F1 complex, beta polypeptide (ATP5B) were calculated according to the standard curve method. Reference genes were selected out of 8 candidate reference genes using the “Genorm” software (Genorm; Primer Design, Southampton, UK). A RT-qPCR program for both RNA copy numbers and host genes of 5 min at 50°C and 20 s at 95°C or direct 3 min at 95°C, followed by 45 cycles of 5 s at 95°C and 30 s at 60°C or 63°C (optimal temperature depending on primers), was performed on a CFX384 Touch™ Real-Time PCR Detection System (Bio-Rad). Primer pairs are listed in Table 1.

**Table 1.**
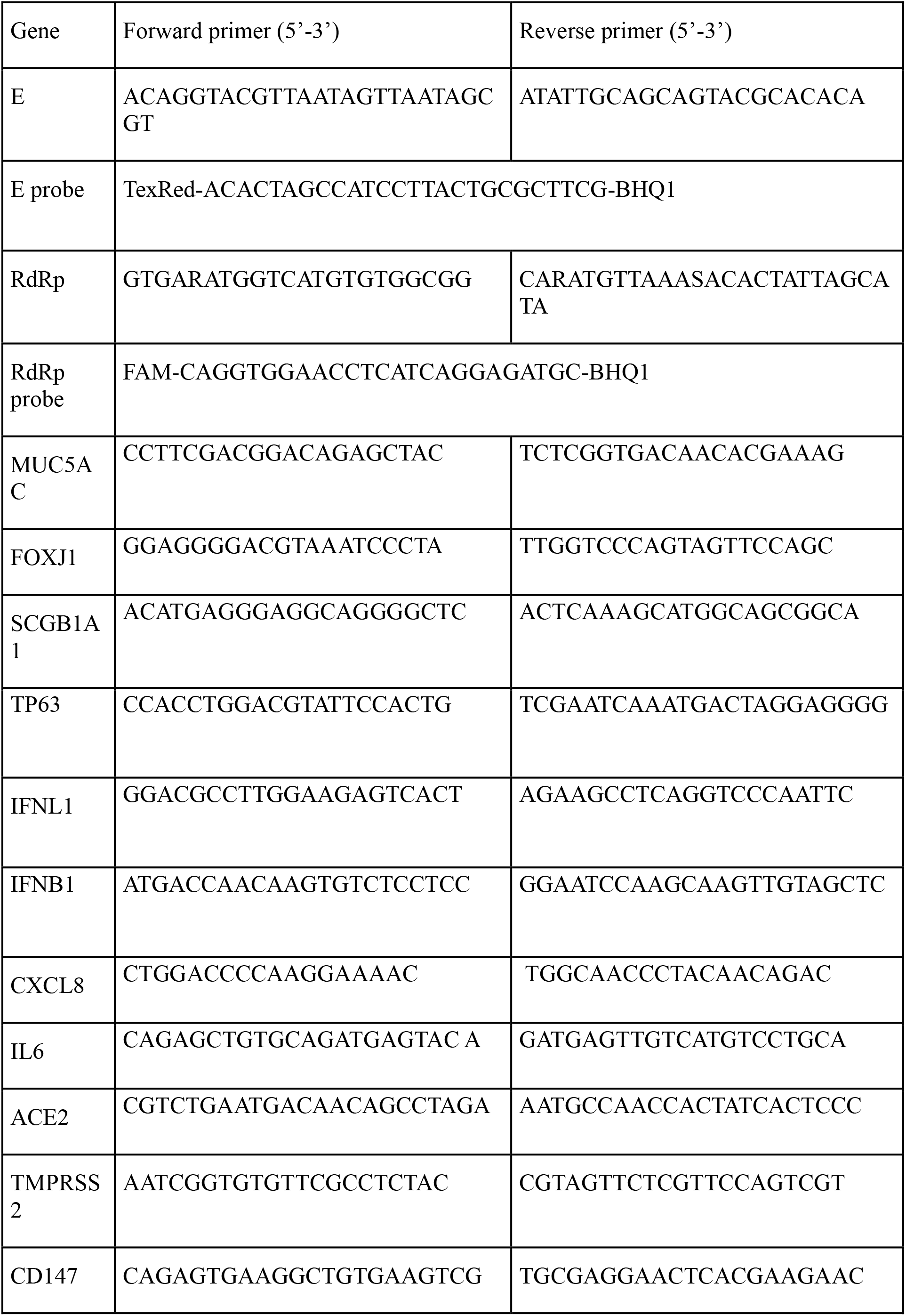
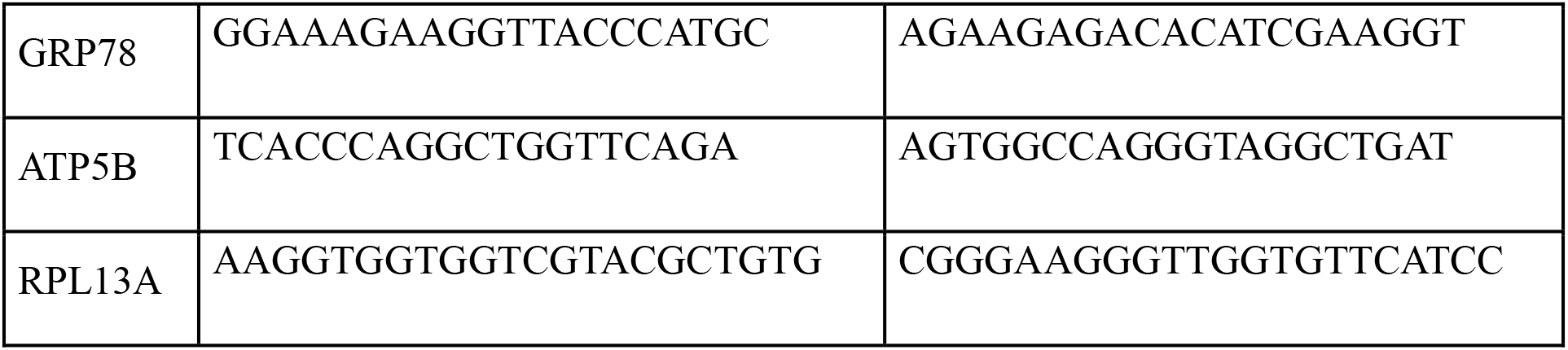
Primer sequences

For quantification of the number of infectious virus particles, the apical wash was serially diluted and infectious titers were determined by plaque assay on Vero E6 as described before (44).

### Immunofluorescence staining

ALI cultures were rinsed using PBS and cells were fixed by adding 3% (w/v) paraformaldehyde diluted in PBS into the basal and apical compartments followed by incubation at room temperature for at least 35 min. Next, inserts were washed two times with PBS and stored in PBS with 10 mM glycine at 4°C until further use. Ice-cold methanol was added for 10 min at 4°C, and PBS containing 1% (w/v) BSA, 0.3% (w/v) Triton-X-100 (PBT) was used to block non-specific binding sites and permeabilize cells for 30 min at 4°C. Membranes were excised from the insert and cut into 4 pieces that were incubated overnight at 4°C with specific antibodies at the following dilutions: rabbit anti-SARS-CoV-2 N antibody (JUC3,1:500, (46)), mouse anti-MUC5AC antibody (1:200; Thermo Fisher Scientific), mouse anti-acetylated α-tubulin (1/100; Sigma Aldrich) or goat anti-FOXJ1 antibody (1:200; R&D, Minneapolis, MN, USA). After washing, membranes were incubated with corresponding secondary antibodies: donkey anti-rabbit, donkey anti-mouse or donkey anti-goat Alexa-fluor antibodies (all diluted 1:200, Thermo Fisher Scientific) and 4’,6-diamidino-2-phenylindole (DAPI, 1:200, Sigma-Aldrich) in the dark for 30 min at room temperature. Next, membranes were transferred to glass slides and covered with prolong gold anti-fading reagent (Thermo Fisher Scientific) and a coverslip (VWR, Amsterdam, the Netherlands). Slides were viewed using a Leica TCS SP8 confocal microscope (Leica Microsystems, Wetzlar, Germany) at 100 x /400 x / 630 x original magnification according to experimental requirements.

## Acknowledgements

This study was supported by a COVID-19 MKMD grant from the Netherlands Organisation for Health Research and Development (ZonMw) and the Dutch Society for the Replacement of Animal Testing (Stichting Proefdiervrij) (grant #114025007). C.S.-B. was supported by the Coordination for the Improvement of Higher Education Personnel (CAPES) (process no. 88881.171440/2018-01), Ministry of Education, Brazil. Part of this research was supported by the Leiden University Fund (LUF), the Bontius Foundation, and donations from the crowdfunding initiative “wake up to corona”. Collection of primary human tracheal epithelial cells was supported by grants from the Deutsche Forschungsgemeinschaft (DFG) (Ta 275/7-1 and Ta 275/8-1) to C.T.

## Supplementary figures

**Supplementary Figure 1. Effect of culture duration on SARS-CoV-2 viral load in ALI-PTEC and ALI-PBEC cultures.** ALI-PBEC/ALI-PTEC (mix of 5 donors) cultured for 3-5 weeks at ALI were infected with SARS-CoV-2 (30,000 PFU per insert). (A) Extracellular viral RNA copies in the apical wash were measured by RT-qPCR. (B) Plaque assay was performed to quantify infectious virus titers in ALI-PBEC or ALI-PTEC. Data are shown as mean ± SEM. Three different donor mixes (1–3) from ALI-PBEC/PTEC were included, each containing biological duplicates.

**Supplementary Figure 2. Cell types infected by SARS-CoV-2 in ALI-PTEC.** Cells (mix of 5 donors) cultured for 5 weeks were infected with SARS-CoV-2 (30,000 PFU per insert). At 72 hpi, cells were fixed and stained with antibodies against acetylated α-tubulin (ciliated cell marker, A) or MUC5AC (goblet cell marker, B) in combination with anti-SARS-CoV-2 N protein antibody (JUC3) and DAPI for nuclear staining. Immunofluorescence images shown are representative for results of 3 independent experiments with 630 x original magnification. White arrows indicate cells that show co-staining of cell markers and viral N protein.

**Supplementary Figure 3. Effect of culture duration on expression of epithelial cell-specific genes by PTEC and PBEC cultures.** ALI-PTEC/PBEC (mix of 5 donors) were differentiated at ALI for 2, 3, 4 or 5 weeks, and analyzed by RT-qPCR to measure gene expression of *SCGB1A1* (club cell marker) and *TP63* (basal cell marker). n=2 independent experiments at week 2 and n=3 independent experiments at 3-5 weeks derived from three donor mixes the same as infection cultures. Data are mean ± SEM. Analysis of differences was conducted using two-way ANOVA with a Tukey/Bonferroni post-hoc test. Significant differences are indicated by *P<0.05.

**Supplementary Figure 4. Effect of long-term DAPT/IL-13 exposure on epithelial cell composition and effect of acute DAPT/IL-13 exposure on susceptibility to SARS-CoV-2 infection.** ALI-PBEC (mix of 4-5 donors) were differentiated for 3 weeks, before addition of DAPT (5 µM) or IL-13 (1 ng/ml), followed by differentiation for an additional 2 weeks. (A) After 5 weeks of differentiation, ALI-PBEC were fixed, stained and analyzed by immunofluorescence using primary antibodies against MUC5AC and FOXJ1 (goblet cell marker, ciliated cell marker) or acetylated α-tubulin together with FOXJ1 (both ciliated cell markers) in combination with DAPI for nuclear staining. The quantification of FOXJ1-positive cells and MUC5AC-positive cells was done by ImageJ software. Data are mean ± SEM. (B) mRNA levels of *FOXJ1* and *MUC5AC* were measured by RT-qPCR. Data are mean ± SEM. Analysis of differences was conducted using paired t test. (C) Short-term treatment with DAPT was performed in cells after 5 weeks of differentiation. Cells were either pre-treated with DAPT for 24 h (Pre-treatment) or post-treated directly after infection (Post-treatment). Intracellular SARS-CoV-2 RNA copies were measured by RT-qPCR and plaque assay was performed with apical washes to quantify infectious virus titers. n=3 independent experiments. Data are mean ± SEM. Fold change in intracellular RNA copies was compared to untreated controls. Statistical analysis was performed using a paired t test. Significant differences are indicated by *P<0.05.

